# Odor-background segregation of unknown odorants based on stimulus onset asynchrony in honey bees

**DOI:** 10.1101/602045

**Authors:** Aarti Sehdev, Paul Szyszka

## Abstract

Animals use olfaction to search for distant objects. Unlike vision, where objects are spaced out, olfactory information mixes when it reaches olfactory organs. Therefore, efficient olfactory search requires segregating odors that are mixed with background odors. Animals can segregate known target odors by detecting short differences in the arrival of odorants from different sources (stimulus onset asynchrony). However, it is unclear whether animals can also use stimulus onset asynchrony to segregate previously unknown odorants that have no innate or learned relevance. Using behavioral experiments in honey bees, we here show that stimulus onset asynchrony also improves odor-background segregation of unknown odorants. The stimulus onset asynchrony necessary to behaviorally segregate unknown odorants is in the range of seconds, which is two orders of magnitude larger than the previously reported stimulus asynchrony sufficient for segregating known odorants. We propose that for unknown odorants, odor-background segregation requires sensory adaptation to the background stimulus.

## INTRODUCTION

Natural olfactory stimuli often comprise several chemically different odorants from different sources that mix together in odor plumes (Celani et al., 2014; Murlis et al., 1992; Riffell et al., 2014). Previous studies suggested that animals perceive odorant mixtures synthetically, that is, they perceive a mixture as a perceptual unit rather than as a list of individual odorants (humans: (Jinks and Laing, 1999); squirrel monkeys: (Laska and Hudson, 1993); rats: (Staubli et al., 1987); spiny lobsters: (Lynn et al., 1994), honey bees: (Chandra and Smith, 1998; Deisig et al., 2001; Smith, 1998)). But, animals often need to segregate information about odorants coming from one source from background odors created by other sources (Hopfield, 1991; Stevenson and Wilson, 2007). Odor plumes are turbulent, which creates filaments of odorant interspersed with non-odorized air (Murlis et al., 2000). Due to this turbulent structure of plumes, odorants that originate from the same source fluctuate with stable relative concentration proportions, forming a homogeneous stream of plumes, whereas odorants from different sources will have varying relative concentration proportions, creating a heterogeneous stream of plumes (Celani et al., 2014). Therefore, homogeneous and heterogeneous plumes could provide the animal with information about the number of odor sources and which odorants belong to the same odor source. In particular, odorants that arise from a single source would arrive at the olfactory organ synchronously, whereas odorants that arise from multiple sources would differ in their arrival times (Erskine et al., 2019; Hopfield, 1991). Accordingly, invertebrates can use both spatial and temporal information from odor plumes for odor-background segregation (spatial: (Andersson et al., 2011; Baker et al., 1998; Hopfield and Gelperin, 1989; Weissburg et al., 2012); temporal: (Saha et al., 2013; Sehdev et al., 2019; Szyszka et al., 2012). Remarkably, tobacco hawk moths can segregate odorant sources separated by only 1 millimeter (Baker et al., 1998), and honey bees can use odorant onset asynchronies as short as 6 milliseconds to segregate a known target odorant (odorant with innate or learned valence) from a background odorant (Szyszka et al., 2012).

All previous studies that have investigated odor segregation based on temporal stimulus cues have in common that the target odorant was known to the animal, either through having an innate valence (Andersson et al., 2011; Baker et al., 1998; Nikonov and Leal, 2002; Weissburg et al., 2012) or via appetitive conditioning (Hopfield and Gelperin, 1989; Saha et al., 2013; Sehdev et al., 2019; Szyszka et al., 2012). It is currently unknown whether animals can use temporal stimulus cues to segregate an unknown odorant (odorant without innate or learned valence) from a background without ever encountering the target odorant alone.

To test whether stimulus onset asynchrony improves segregating an unknown odorant from a background odor, we trained honey bees in a classical conditioning assay (Bitterman et al., 1983) to associate a target odorant A mixed to a background B, with a sucrose reward. The background odor B started either before A (asynchronous mixture) or simultaneously with A (synchronous mixture). Thus, during conditioning, bees could never encounter A alone. We then tested whether bees had segregated A from the background B during conditioning by testing their response to A or to a novel odorant. We found that bees were able to segregate A from both synchronous and asynchronous mixtures with the background B, but bees performed better when the background B started 20 or 5 seconds before A, as compared to when B started between 1 second before to 0.2 seconds after the onset of A. These data suggest that bees can use stimulus onset asynchrony to segregate an unknown odorant from an olfactory background, but not on the millisecond timescale seen for known target odorants.

## MATERIALS AND METHODS

### Animals

Worker honey bees (*Apis mellifera*) were collected from the entrance of outdoor hives at University of Konstanz between 09:00 and 12:00 between June 2016 and December 2016. Bees were anaesthetized using ice and fixed into a holder using sticky tape, so that proboscis, antennae and mandibles were freely movable. The fixed bees were left undisturbed and starved for three hours before the conditioning procedure started, in order to encourage a response to a sucrose reward. We only used bees that showed responses to sucrose when presented at the antennae. Up to 16 bees were conditioned in parallel in each experimental session.

### Odorant delivery

Bees were conditioned to a target odorant (A) of either 1-hexanol or nonanal. These odorants were chosen based on their perceived dissimilarity to each other (Guerrieri et al., 2005). To make the background segregation task challenging, we presented a complex background mixture (B), comprised of four odorants: 1-octanol, heptanal, hexanal and 2-hexanone. These odorants were also chosen based on their perceived dissimilarity to the target odorants (Guerrieri et al., 2005). All pure odorants were supplied by Sigma-Aldrich.

Odors were delivered to the antennae of the bee using a custom-made olfactory stimulator, as designed and described in Raiser, Galizia and Szyszka (2017). Pure odorants were kept in 20 ml glass vials (Schmidlin) sealed with a Teflon septum; the headspace of odorized air was extracted and drawn into the air dilution system using flowmeters (112-02GL, Analyt-MTC) and an electronic pressure control (35898; Analyt-MTC). The cross-sectional area of all odor vials was the same, so that the odorants would have the same concentration when using the same airflow. The olfactory stimulator used three channels: one for nonanal, one for 1-hexanol and one for the background mixture. To create the background odor, the four odorants were mixed by connecting the four odorant vials in series and passing 50 ml/min of air through their headspaces allowing the four odorants to mix. For each odor channel, the rate of air flow was 300 ml/min, with 50 ml/min of odorant combined with a dilution of 250 ml/min of clean air. The total air flow at the outlet of the stimulator was 4.7 L/min, with an airspeed of 1 m/s. The outlet of the stimulator had an inner diameter of 1 cm and was positioned 1 cm in front of the center of the bees’ head.

The valves of the olfactory stimulator were controlled by a compact RIO system equipped with a digital I/O module NI-9403 (National Instruments) using software written by Stefanie Neupert in LabVIEW 2011 SP1 (National Instruments). The odorant vials were constantly flushed with air throughout the experiment, so that the headspace concentration reached a dynamic steady state. To generate the synchronous odorant mixture, two odor channels (either channels 1 and 3 or channels 2 and 3) were opened simultaneously with a single trigger. To generate the asynchronous mixtures, the two odor channels were opened using two triggers with a time delay between them. All odorants were removed swiftly from the setup area via an exhaust placed behind the bee.

### Conditioning paradigm

All experiments are based on classical absolute conditioning by pairing an olfactory stimulus (conditioned stimulus) with a sucrose reward (unconditioned stimulus) (Bitterman et al., 1983). The sucrose reward was 1.25 Mol sucrose-water solution and was applied by a metal pinhead (1 mm diameter) that was dipped into the sucrose. The sucrose reward was presented for four seconds, first to the antennae to induce the proboscis extension response and then to the proboscis to allow for feeding. Bees were conditioned for five trials and the inter-stimulus interval was approximately 14 minutes. The time between the final conditioning trial and the test was 30 minutes. Overall, 522 bees were tested.

During conditioning, fixed bees were placed in front of the stimulator, where they were left to acclimatize to the airflow for 20 seconds. After this, the valves for the background B were opened; after a delay of either 20 s (B20A), 5 s (B5A), 1 s (B1A) or 0.2 s (B0.2A), the target odorant A was released. For the synchronous mixture AB, the valves for the background B and the target odorant A were opened simultaneously. Three seconds after the onset of A, the sucrose reward was presented. LEDs were used to indicate to the experimenter when to present sucrose to the bees and when the different odorants were released from the stimulator valves.

All experiments performed were balanced, so that odorant A was either 1-hexanol or nonanal. During the test, bees were presented with the target odorant A (either 1-hexanol or nonanal) and a novel odorant N (either nonanal or 1-hexanol) over two consecutive trials in a balanced order. The background odorant B was not used during testing.

### Quantifying conditioned response

We monitored the conditioned response as the occurrence of the proboscis extension reflex during the odorant stimulation. We counted a proboscis extension reflex only when the proboscis was not extended before odorant onset and when it was extended horizontally during the odorant stimulation not overlapping with the sucrose reward. The conditioned response was documented in a binary form. For each conditioning trial, the percentage of bees that showed proboscis extension reflex to each odorant was recorded. For those treatments where the arrival of the background mixture could be discerned visually from the arrival of target odorant A by the experimenter (B20A, B5A), the proboscis extension reflex was recorded for both A and B. During the test, the presence or absence of proboscis extension reflex to the conditioned odorant was recorded as a 1 or 0 respectively. To assess the associative memory performance, we separated responses into “correct” and “incorrect” responses. Bees that responded to the A during the test but not the novel odorant N were given a score of 1; all other responses were deemed incorrect and given a score of 0. The proportion of correct responses was then statistically compared between groups.

Bees that died during the experiment, or those that did not respond to sucrose when delivered to the antennae at the end of the experiment, were discarded from the analysis.

### Statistical analysis

For all data analysis, R version 3.5.2 was used (R Core Team, 2012). All statistical tests were performed using Bayesian data analysis, based on Korner-Nievergelt *et al.* (2015).

To investigate the effect of the target odorant A and novel odorant N on conditioned response, we used a binomial generalized linear model, with conditioned response as the binary response variable (1 = conditioned response, 0 = no conditioned response). We used the logistic regression (logit) link function. The target odorant A and novel odorant N were used as explanatory variables. We used an improper prior distribution (flat prior) and simulated 100 000 values from the posterior distribution of the model parameters using the function “sim” from the package “arm”. The means of the simulated values from the posterior distributions of the model parameters were used as estimates, and the 2.5 % and 97.5 % quantiles as the lower and upper limits of the 95 % credible intervals. To test for differences between conditioned responses to the target odorant A and novel odorant N, we compared the probabilities of conditioned response by calculating the proportion of simulated values from the posterior distribution that were larger in the target odorant A than in the novel odorant N. A posterior probability of, for example, 0.953 for the comparison between the target odorant A and the novel odorant N (*p(A > N)* = 0.953) means that one can be 95% certain that the probability of conditioned response is greater for A than for N. We declared an effect to be significant if the proportion was greater than 0.95.

To investigate the effect of mixtures and single odorants or of synchronous and asynchronous mixtures on correct responses, we used the same analysis as above, with the appropriate explanatory variables.

## RESULTS

To investigate bees’ capability to segregate an unknown odorant from a background, we conditioned fixed bees to associate a target odorant A (either 1-hexanol or nonanal), which was mixed into an olfactory background B (mixture of 1-octanol, heptanal, hexanal and 2-hexanone), with a sucrose reward. Sucrose was always presented 3 seconds after the onset of A. 30 minutes after the last conditioning trial, the conditioned response to A alone or to a novel odorant (N) (either 1-hexanol or nonanal) was tested in each bee. To eliminate between-session variability, all data shown in a given panel of a figure were collected in parallel during the same experimental sessions. Accordingly, data points should be compared within panels, but not between panels (original data are available in Data S1).

### Stimulus onset asynchrony between mixed odorants impairs recognition of the components

We firstly wanted to confirm the results from previous studies that mixing odorants impairs bees’ recognition of the individual odorants (Chandra and Smith, 1998; Deisig et al., 2001; Smith, 1998). We conditioned bees to associate a synchronous mixture of odorants A and B (AB, where A and B start at the same time) with a sucrose reward (Fig. 1A). A second group was conditioned to A without B (Fig. 1A).

**Figure 1.**
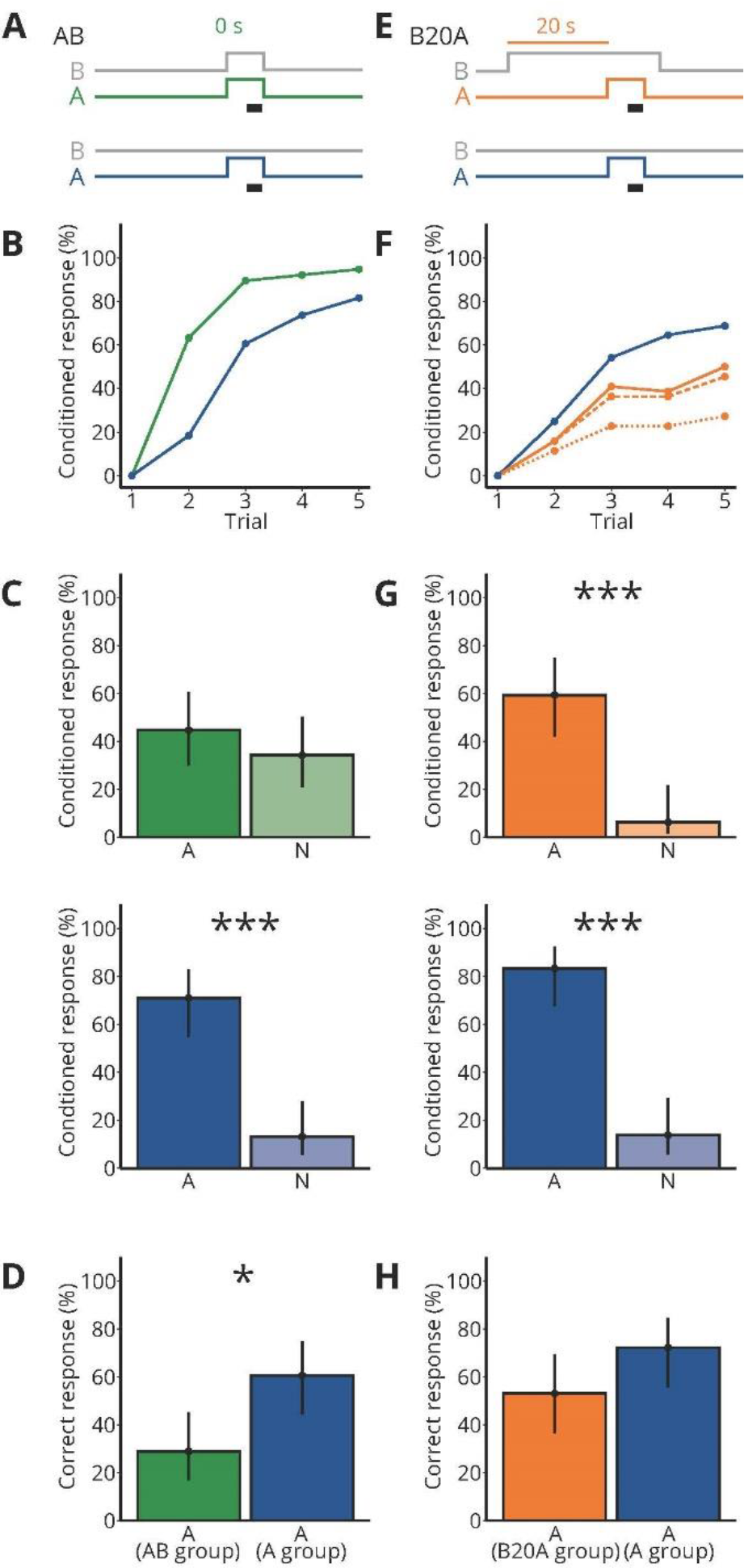
Mixing an unknown odorant A with a background odor B impairs recognition of A. **(A)** Valve states for creating odorant pulses for the synchronous mixture AB and the control A. For AB, both the target A and the background B (grey) were turned on synchronously and were presented for 7 seconds. The black bar indicates the 4 seconds when the sucrose reward was given. For the control A, the target A (blue) was presented for 7 s. B (grey) was not presented. **(B)** Each bee received 5 rewarded conditioned trials either with AB (green) or A (blue); the percentage of bees responding to the odorants is shown. N = 38 bees conditioned to AB and 38 bees conditioned to A. **(C)** During the two test trials, each bee was stimulated with A and a novel odorant N. Percentage of bees responding to A and to N for bees conditioned to AB (green) and bees conditioned to A (blue). Points represent means and vertical lines represent 95 % credible intervals for all panels in this figure. Stars indicate significant differences between means for all panels in this figure (*: probability for a difference between both means p > 0.95; ***: p >0.999). **(D)** Percentage of correctly responding bees during the test (response to A but not to N) for bees conditioned to AB (green) and A (blue). **(E)** Valve states for creating odorant pulses for the asynchronous mixture B20A and the control A. For B20A, the target A (orange) was turned on 20 seconds after the background B (grey). A was presented for 7 seconds. B ended 3 seconds after A ended. Same control as in (A). **(F)** Same as in (B), but for B20A. Percentage of bees responding to only the target A (dotted line), only the target A or to both A and B within the same trial (dashed line), and to A and/or B (solid line) for bees conditioned to B20A (orange) and A alone (blue). N = 32 bees conditioned to B20A and 36 bees conditioned to A. **(G)** Same as in (C), but for B20A. **(H)** Same as in (D), but for B20A.

By the fifth conditioning trial, 95 % of bees conditioned to AB responded and 82 % of the bees conditioned to A alone responded (Fig. 1B). To test whether the bees associated odorant A with the sucrose reward during conditioning, we then recorded the response to A or to a novel odorant N in the absence of sucrose (the order of A and N was balanced across bees) (Fig. 1C). Significantly more bees responded to A than to N in the group conditioned to A (*p(A > N)* > 0.999), but not in the group conditioned to AB (*p(A > N)* = 0.826), indicating that bees recognize A as a stimulus predictive for the sucrose reward when conditioned to A but not when conditioned to AB. During the test, bees could respond in several ways. They could respond correctly, that is showing a conditioned response to A and not to N; alternatively they could generalize by responding to both odorants, or show a lack of response to A. To determine whether the groups conditioned to the asynchronous or synchronous background differed in their expression of correct responses, we calculated the percentage of bees’ correct responses in the test (Fig. 1D). The percentage of correctly responding bees was lower when conditioned to the synchronous mixture AB than with A alone (*p(A(A group) > A(AB group)* = 0.997), confirming earlier studies showing that mixing odorants impairs the recognition of individual odorant components, indicating that the perception of odorant mixture is partly synthetic (Chandra and Smith, 1998; Deisig et al., 2001; Smith, 1998).

Because stimulus onset asynchrony can improve odor-background segregation for known target odorants (Andersson et al., 2011; Baker et al., 1998; Hopfield and Gelperin, 1989; Sehdev et al., 2019; Szyszka et al., 2012), we asked whether mixing A with an asynchronous background would still impair the recognition of A in a mixture with B. We presented B for 30 s and added a 7 s long target odorant A after 20 s (B20A; Fig. 1E). B always stopped 3 s after A stopped to make sure that bees would never encounter A without B. The sucrose reward was presented 4 s after the onset of A (Fig. 1E). As a control, bees were conditioned to A without any background. By the fifth conditioning trial, 27 % of the bees conditioned to B20A responded to A only and 45 % responded to both A and B, and 50 % responded to A and/or B, and 69 % of the bees conditioned to A alone responded to A (Fig. 1F).

During the test, more bees conditioned to B20A responded to A than to N (*p(A > N)* > 0.999), as did bees conditioned to A (*p(A > N)* > 0.999) (Fig. 1G), showing that bees could segregate A from B20A during the conditioning. Moreover, the percentage of correctly responding bees did not significantly differ when conditioned to B20A than to A alone (*p(A(A group) > A(B20A group))* = 0.947) (Fig. 1H). These results suggest that bees could segregate a target odorant A better in B20A than in AB.

### Long onset asynchrony improves odor-background segregation of an unknown odorant

To determine the limit of stimulus onset asynchrony that bees could use for odor-background segregation, we investigated three onset asynchronies between the background B and a following target odorant A: 5 s (B5A), 1 s (B1A) and 0.2 s (B0.2A) (Fig. 2A). We compared bees’ capability to segregate the target odorant A against a parallel group of bees that was conditioned to the synchronous mixture AB (A and B had a synchronous onset, but different to the experiment in Fig. 1A, B stopped 3 s after A stopped). During the fifth conditioning trial, 36 % of bees conditioned to B5A responded to A only, 55 % responded to A and B, and 98 % responded to A and/or B (Fig. 2B). In comparison, 93 % of bees conditioned to AB responded to AB (Fig. 2B).

**Figure 2.**
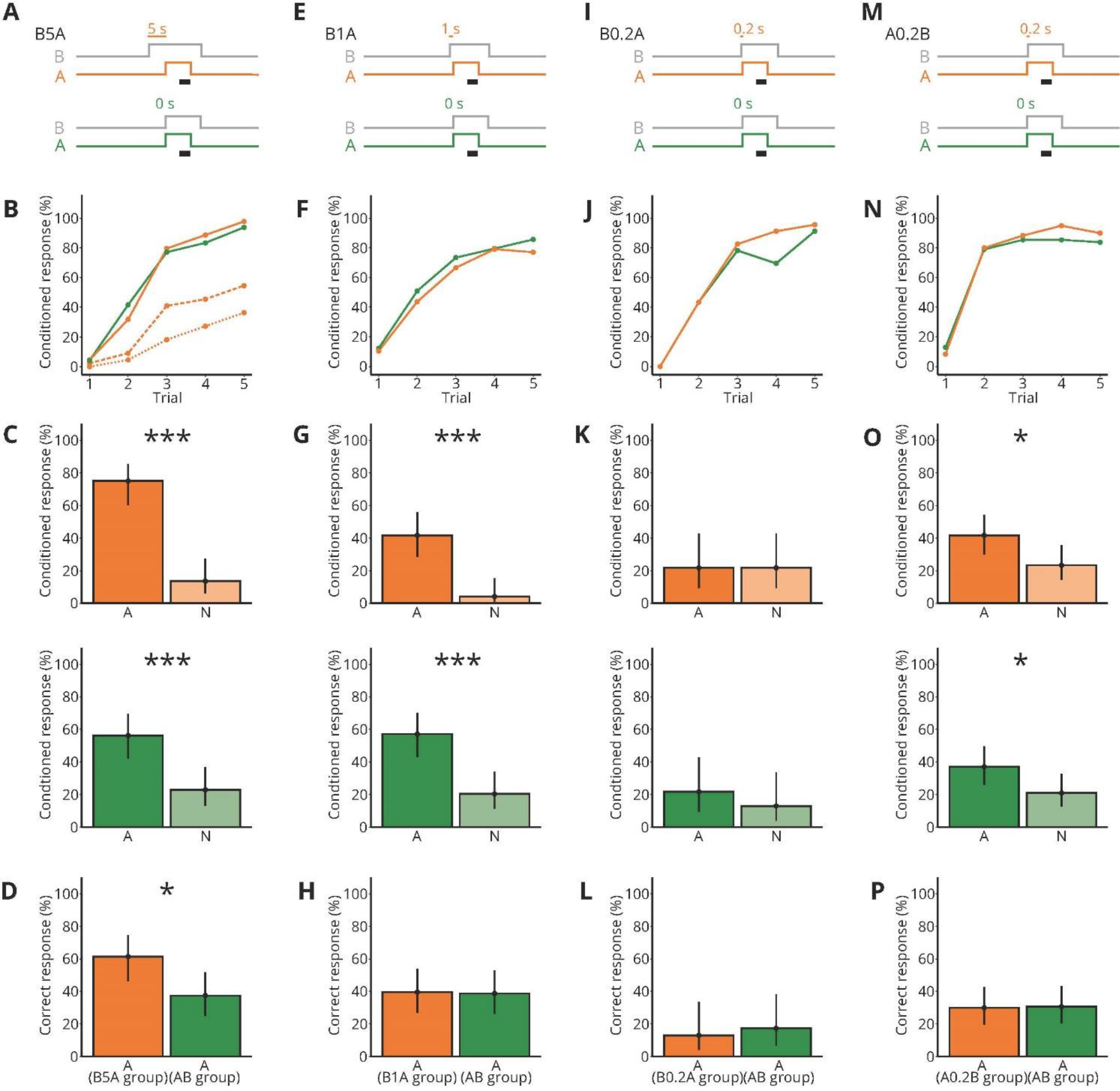
Five seconds onset asynchrony between a background B and a following unknown odorant A improves odor-background segregation. **(A)** Valve states for creating odorant pulses for the asynchronous mixture B5A and the synchronous mixture AB. For B5A, the target A turned on 5 seconds after B (grey). A was presented for 7 seconds. B ended 3 seconds after A ended. The black bar indicates the 4 seconds when the sucrose reward was given. For AB, both the target A and the background B (grey) were turned on synchronously. **(B)** Each bee received 5 rewarded training trials either with B5A (orange) or AB (green). Percentage of bees responding to only the target A (dotted line), only the target A or to both A and B within the same trial (dashed line), and to A and/or B (solid line). N = 44 bees conditioned to B5A and 48 bees conditioned to AB. **(C)** During the two test trials, bees were stimulated with A or a novel odorant N. Percentage of response to A and to N for bees conditioned to B5A (orange) and bees conditioned to AB (green). Points represent means and vertical lines represent 95 % credible intervals for all panels in this figure. Stars indicate significant differences between means for all panels in this figure (*: probability for a difference between both means p > 0.95; ***: p >0.999). **(D)** Percentage of correctly responding bees during the test (response to A but not to N) for bees conditioned to B5A (orange) and A (green). **(E)** Same as in (A), but for B1A. The target A turned on 1 second after B. **(F)** Same as in (B), but for B1A as asynchronous mixture. Separate responses to A and B were not distinguishable. N = 48 bees conditioned to B1A and 49 bees conditioned to AB. **(G)** Same as in (C), but for B1A. **(H)** Same as in (D), but for B1A. **(I)** Same as in (A), but for B0.2A.The target A turned on 0.2 seconds after B. **(J)** Same as in (F), but for B0.2A as asynchronous mixture. N = 23 bees conditioned to B0.2A and 23 bees conditioned to AB. **(K)** Same as in (C), but for B0.2A. **(L)** Same as in (D), but for B0.2A. **(M)** Same as in (A), but for A0.2B. The target A turned on 0.2 seconds before B. **(N)** Same as in (F), but for A0.2B as asynchronous mixture. N = 60 bees conditioned to A0.2B and 62 bees conditioned to AB. **(O)** Same as in (C), but for A0.2B. **(P)** Same as in (D), but for A0.2B.

The bees conditioned to B5A responded more to A than to N during the test (*p(A > N)* > 0.999), showing that they were able to segregate A from the background B during conditioning (Fig 2C). The AB-conditioned bees also responded more to A than to N (*p(A > N)* = 0.999) (Fig. 2C), showing that they also could segregate A from B. However, bees conditioned to B5A showed more correct responses than bees conditioned to AB (*p(A(B5A group) > A(AB group))* = 0.989) (Fig. 2D). This shows that stimulus onset asynchrony of 5 seconds improves segregating a target odorant A from an asynchronous background (B5A) as compared to synchronous presentation of both odorants (AB).

### One second onset asynchrony or shorter does not improve odor-background segregation

Bees conditioned to the asynchronous mixtures B1A and B0.2A (Fig. 2 E and I) showed similar responses to the synchronous mixtures AB during conditioning. The onset asynchrony between background B and target A of 1 second or less was too short to determine whether their conditioned response was to A or to B. During the fifth conditioning trial, 77 % and 86 % responded to B1A and AB, respectively (Fig. 2F), and 96 % and 91% responded to B0.2A and AB, respectively (Fig. 2J).

During the test, bees of the B1A group and the parallel AB control group showed a higher percentage of responses to A than to N (group B1A: *p(A > N)* > 0.999; ; group AB: *p(A > N)* > 0.999; Fig. 2G). However, bees of the B0.2A group and the parallel AB control group showed no difference in their responses to A and N (group B0.2A *p(A > N)* = 0.501; group AB: *p(A > N)* = 0.781; Fig. 2K), indicating that neither bees of the B0.2A group nor of the parallel AB control group segregated A during conditioning. The differences in bees capability to segregate A from the AB control during conditioning between the experimental sessions shown in Fig. 2G and 2K indicates that bees’ capability to segregate an odorant from a mixture may vary and depend on factors other than the experimentally controlled factors (e.g., season, bees’ age). For both the B1A group and the B0.2A group, there was no difference in the proportion of correct responses to the AB-conditioned control for bees (*p(A(B1A group) > A(AB group))* = 0.532; Fig. 2H; *p(A(B0.2A group) > A(AB group))* = 0.343; Fig. 2L).

In summary, bees were able to segregate A from B when conditioned to an asynchronous or synchronous mixture (except for B0.2A). However bees showed a higher percentage of correct responses for an onset asynchrony of 5 seconds between the background B and target A, and not for shorter onset asynchronies. Although the AB-conditioned controls were also able to segregate A, the proportion of correct responses in the test was either lower than the proportion of correct responses from bees conditioned to the asynchronous mixture (B5A), or there was no difference between the proportion of correct responses in the test between bees conditioned to AB or to the asynchronous mixtures (B1A, B0.2A).

In all of the previous experiments, during conditioning, B was presented before A, thus bees could never experience A alone. We therefore asked whether presenting A before B would improve segregation of A from B. We conditioned bees using an asynchronous mixture of A and B, in which B started 0.2 seconds after the onset of A (A0.2B) (Fig. 2M). Thus bees experienced the target odorant A alone for 0.2 seconds before the onset of the background B. Again, we used the synchronous mixture AB as a control. During the fifth conditioning trial 90 % of the bees responded to A0.2B and 83 % responded to AB (Fig. 2N). In the test, bees conditioned to A0.2B showed a higher percentage of conditioned responses to A than to N (*p(A > N)* = 0.983), as did the bees conditioned to AB (*p(A > N)* = 0.975) (Fig. 2O). However the percentage of correct responses during the test was not different between the two groups (*p(A(A0.2B group) > A(AB group))* = 0.471) (Fig. 2M). Therefore, even encountering the target odorant A alone for a short time does not improve its segregation from a mixture.

## DISCUSSION

We asked whether honey bees can use stimulus onset asynchrony to segregate an unknown odorant A (odorant without innate or learned valence) from an olfactory background B. We found that stimulus onset asynchronies of 5 seconds or more improved odor-background segregation, while stimulus onset asynchronies of 1 seconds or less did not. This timescale for segregating unknown odorants based on stimulus onset asynchrony is at least two orders of magnitude slower than for segregating known odorants (Baker et al., 1998; Sehdev et al., 2019; Szyszka et al., 2012). These results suggest that the neural mechanisms for odor-background segregation based on stimulus onset asynchrony differ for known and unknown odorants.

### Perception of odorant mixtures

The finding that bees were impaired in recognizing the odorant A in a mixture with B (Fig. 1A-D) is in line with previous studies which suggested that honey bees perceive odorant mixtures partly synthetically (Chandra and Smith, 1998; Deisig et al., 2001, 2003; Müller et al., 2000; Smith, 1998). However, there is also evidence for analytic mixture perception in bees: when bees are conditioned to a mixture and afterwards are tested with the single odorants, they respond to most of the single odorants (Laloi *et al.*, 1999; Reinhard *et al.*, 2010). Analytic mixture perception has also been demonstrated in blocking experiments, in which previous conditioning to odorant A reduces conditioning to B during conditioning with AB, because A already predicts the reward (Smith and Cobey, 1994) (but see (Gerber and Ullrich, 1999) for an opposing view).

The impairment of bees’ capability to segregate an odorant A from a mixture with B could reflect an impaired detection of odorant A due to synthetic mixture perception, or it could reflect an impaired learning of odorant A due to overshadowing (Pavlov et al., 1927). In honey bees, the potential of one odorant to overshadow another odorant increases with its concentration (Pelz et al., 1997; Reinhard et al., 2010). In our setup, the concentration of B likely was higher than the concentration of A because we used a mixture of 4 odorants as background B and only one odorant as A (see Methods), and the vapor pressures of the B odorants were up to 1 order of magnitude higher than those of the A odorants (A odorants: 1-hexanol: 0.1 kPa, nonanal: 0.05 kPa; B odorants: 1-octanol: 0.001 kPa, heptanal: 0.5 kPa, hexanal: 1.5 kPa, 2-hexanone: 1.5 kPa; all at 25°C). It is therefore plausible to assume that B overshadowed A and was learned better than A.

### Odor-background segregation based on relational stimuli

Segregating an unknown odorant from a mixture is a blind source separation problem which requires more information than just the chemical odorant identity (Hendin et al., 1994). The physics of odorant dispersion adds relational information to the chemical odorant identity, as odorants from the same source form plumes with relatively stable odorant concentration proportions (homogeneous plumes), while odorants from different sources form plumes with variable odorant concentration proportions (heterogeneous plumes) (Celani et al., 2014; Hopfield, 1991). Indeed, animals can use these relational stimuli to detect whether odorants originate from the same or different sources. For example, honey bees can segregate an unknown odorant A from a mixture ABC when the concentrations of B and C vary from trial to trial (Wright and Smith, 2004). Similarly, mice can segregate an unknown odorant from a multi-odorant background whose composition varies from trial to trial (Rokni et al., 2014).

Besides this trial-to-trial variability of odorant concentration or composition, animals can also use stimulus onset asynchrony between mixed odorants for segregating odorants from mixtures (Andersson et al., 2011; Baker et al., 1998; Hopfield and Gelperin, 1989; Saha et al., 2013; Sehdev et al., 2019; Szyszka et al., 2012; Weissburg et al., 2012). However, so far, this capability has only been demonstrated for known odorants that had either an innate or learned valence.

Our data suggest that honey bees can use stimulus onset asynchrony also for segregating unknown odorants. However the required stimulus onset asynchrony for segregating an unknown odorant is in the range of seconds rather than in the range of milliseconds as is the case for known odorants (Baker et al., 1998; Sehdev et al., 2019; Szyszka et al., 2012).

### Neural mechanisms for segregating known versus unknown odorants from background

The longer stimulus onset asynchrony required for segregating an unknown odorant from background suggests a lower temporal resolution for the encoding of unknown odorants as compared to known odorants. The temporal resolution with which unknown odorants can be encoded is limited by the time window over which the olfactory system integrates odor-evoked responses of olfactory receptor neurons (Egea-Weiss et al., 2018; Jeanne and Wilson, 2015). The finding that bees required stimulus onset asynchronies longer than one second for odor-background segregation, indicates bees’ olfactory system needs to integrate olfactory receptor neuron responses for more than one second in order to identify unknown odorants.

In contrast, the finding, that insects can use stimulus onset asynchronies of a few milliseconds for odor-background segregation when the target odorant is known (Baker et al., 1998; Sehdev et al., 2019; Szyszka et al., 2012) indicates that the temporal resolution with which odorants can be encoded is higher for known than for unknown odorants. The temporal resolution for encoding a known odorant could be higher, because a known odorant activates temporally precise pattern recognition neurons that are tuned to the specific neuronal activity pattern evoked by the known odorant. Those pattern recognition neurons could be lateral horn neurons in case of odorants with innate valence (Jeanne et al., 2018; Jefferis et al., 2007; Roussel et al., 2014; Strutz et al., 2014), mushroom body output neurons in case of odorants with learned valence (Aso et al., 2014; Hige et al., 2015; Strube-Bloss et al., 2011) (Aso et al., 2014; Hige et al., 2015; Strube-Bloss et al., 2011), or Kenyon cells in case of previously encountered odorants (Cassenaer and Laurent, 2007; Stopfer and Laurent, 1999). Therefore, whether odorants originate from the same or different sources could be detected by coincidence-detecting neurons that receive input from those pattern recognition neurons (Sehdev et al., 2019).

Our finding that stimulus onset asynchronies of 5 seconds or longer, but not of 1 second or shorter, improve odor-background segregation, is consistent with the hypothesis that for unknown odorants, odor-background segregation depends on sensory adaptation. Sensory adaptation was previously proposed to underlie odor-background segregation in vertebrates (Kadohisa and Wilson, 2006; Linster et al., 2007). Neurons in the piriform cortex (analogous to the mushroom bodies) adapt to a constant odorant stimulus but remain responsive to a novel odorant pulse that arrives a few tens of seconds after the onset of the background odorant (Wilson, 1998).

Similarly, insect olfactory receptor neurons and projection neurons (in *Drosophila*) adapt to a constant odorant stimulus (Bhandawat et al., 2007; Cafaro, 2016; de Bruyne et al., 2001; Martelli et al., 2013; Nagel and Wilson, 2011) but they remain responsive to a novel odorant pulse that arrives a few seconds after the onset of the background odorant (Cafaro, 2016). In honey bees, neural responses start adapting within a few hundreds of milliseconds (projection neurons: (Krofczik et al., 2009; Sachse and Galizia, 2002); Kenyon cells: (Farkhooi et al., 2013; Froese et al., 2014; Szyszka et al., 2005)). Therefore, in honey bees, like in vertebrates, sensory adaptation could filter stable background odors while the neural responses to a novel odorant with an asynchronous onset would not be diminished, thus facilitating the segregation of a novel odorant with a later onset than the background.

## CONFLICT OF INTEREST

The authors declare that the research was conducted in the absence of any commercial or financial relationships that could be construed as a potential conflict of interest.

## AUTHOR CONTRIBUTIONS

A.S. and P.S. conceptualized and designed the study. A.S. performed the data collection and performed the statistical analysis. A.S. and P.S. wrote the manuscript. P.S. supervised the study.

## FUNDING

This project was funded by the Human Frontier Science Program (RGP0053/2015) to PS and by the IMPRS Organismal Biology, University of Konstanz to AS.

## ACKNOWLEDGMENTS

We thank Stefanie Neupert and C. Giovanni Galizia for comments on the manuscript and Doreen Taube and Sabrina Henke for help with the behavioral experiments.

## REFERENCES

Andersson, M. N., Binyameen, M., Sadek, M. M., and Schlyter, F. (2011). Attraction Modulated by Spacing of Pheromone Components and Anti-attractants in a Bark Beetle and a Moth. J. Chem. Ecol. 37, 899–911. doi:10.1007/s10886-011-9995-3.

Aso, Y., Sitaraman, D., Ichinose, T., Kaun, K. R., Vogt, K., Belliart-Guérin, G., et al. (2014). Mushroom body output neurons encode valence and guide memory-based action selection in Drosophila. Elife 3, e04580. doi:10.7554/eLife.04580.

Baker, T. C. T., Fadamiro, H. Y., and Cosse, A. A. A. (1998). Moth uses fine tuning for odour resolution. Nature 393, 530–530. doi:10.1038/31131.

Bhandawat, V., Olsen, S. S. R., Gouwens, N. W. N., Schlief, M. L. M., and Wilson, R. R. I. (2007). Sensory processing in the Drosophila antennal lobe increases reliability and separability of ensemble odor representations. Nat. Neurosci. 10, 1474–1482. doi:10.1038/nn1976.

Bitterman, M. E., Menzel, R., Fietz, A., Schäfer, S., and Schafer, S. (1983). Classical Conditioning of Proboscis Extension in Honeybees (Apis mellifera). J. Comp. Psychol. 97, 107–119. doi:10.1037/0735-7036.97.2.107.

Cafaro, J. (2016). Multiple sites of adaptation lead to contrast encoding in the Drosophila olfactory system. Physiol. Rep. 4, e12762. doi:10.14814/phy2.12762.

Cassenaer, S., and Laurent, G. (2007). Hebbian STDP in mushroom bodies facilitates the synchronous flow of olfactory information in locusts. Nature 448, 709–713. doi:10.1038/nature05973.

Celani, A., Villermaux, E., and Vergassola, M. (2014). Odor landscapes in turbulent environments. Phys. Rev. X 4, 1–17. doi:10.1103/PhysRevX.4.041015.

Chandra, S., and Smith, B. H. (1998). An analysis of synthetic processing of odor mixtures in the honeybee (Apis mellifera). J. Exp. Biol. 201, 3113–3121. Available at: http://jeb.biologists.org/cgi/content/abstract/201/22/3113 [Accessed: August 22, 2018].

de Bruyne, M., Foster, K., and Carlson, J. R. (2001). Odor Coding in the Antenna. Neuron 30, 537–552.

Deisig, N., Lachnit, H., Giurfa, M., and Hellstern, F. (2001). Configural olfactory learning in honeybees: Negative and positive patterning discrimination. Learn. Mem. 8, 70–78. doi:10.1101/lm.8.2.70.

Deisig, N., Lachnit, H., Sandoz, J.-C., Lober, K., and Giurfa, M. (2003). A Modified Version of the Unique Cue Theory Accounts for Olfactory Compound Processing in Honeybees. Learn. Mem. 10, 199–208. doi:10.1101/lm.55803.

Egea-Weiss, A., Renner, A., Kleineidam, C. J., and Szyszka, P. (2018). High Precision of Spike Timing across Olfactory Receptor Neurons Allows Rapid Odor Coding in Drosophila. iScience 4, 76–83. doi:10.1016/j.isci.2018.05.009.

Erskine, A., Ackels, T., Dasgupta, D., Fukunaga, I., and Schaefer, A. T. (2019). Mammalian olfaction is a high temporal bandwidth sense. bioRxiv, 570689. doi:10.1101/570689.

Farkhooi, F., Froese, A., Muller, E., Menzel, R., and Nawrot, M. P. (2013). Cellular Adaptation Facilitates Sparse and Reliable Coding in Sensory Pathways. PLoS Comput. Biol. 9, e1003251. doi:10.1371/journal.pcbi.1003251.

Froese, A., Szyszka, P., and Menzel, R. (2014). Effect of GABAergic inhibition on odorant concentration coding in mushroom body intrinsic neurons of the honeybee. J. Comp. Physiol. A Neuroethol. Sensory, Neural, Behav. Physiol. 200. doi:10.1007/s00359-013-0877-8.

Gerber, B., and Ullrich, J. (1999). No evidence for olfactory blocking in honeybee classical conditioning. J. Exp. Biol. 202 (Pt 13, 1839–54. Available at: http://www.ncbi.nlm.nih.gov/pubmed/10359686.

Guerrieri, F., Schubert, M., Sandoz, J. C., and Giurfa, M. (2005). Perceptual and neural olfactory similarity in honeybees. PLoS Biol. 3, 0718–0732. doi:10.1371/journal.pbio.0030060.

Hendin, O., Horn, D., and Hopfield, J. J. (1994). Decomposition of a mixture of signals in a model of the olfactory bulb. Proc. Natl. Acad. Sci. U. S. A. 91, 5942–6. Available at: http://www.ncbi.nlm.nih.gov/pubmed/8016093 [Accessed: April 7, 2019].

Hige, T., Aso, Y., Modi, M. N., Rubin, G. M., and Turner, G. C. (2015). Heterosynaptic Plasticity Underlies Aversive Olfactory Learning in Drosophila. Neuron 88, 985–998. doi:10.1016/j.neuron.2015.11.003.

Hopfield, J. F., and Gelperin, A. (1989). Differential conditioning to a compound stimulus and its components in the terrestrial mollusc Limax maximus. Behav. Neurosci. 103, 329–333. doi:10.1037/0735-7044.103.2.329.

Hopfield, J. J. (1991). Olfactory computation and object perception. Proc. Natl. Acad. Sci. 88, 6462–6466. doi:10.1073/pnas.88.15.6462.

Jeanne, J. M., Fişek, M., and Wilson, R. I. (2018). The Organization of Projections from Olfactory Glomeruli onto Higher-Order Neurons. Neuron 98, 1198–1213.e6. doi:10.1016/j.neuron.2018.05.011.

Jeanne, J. M., and Wilson, R. I. (2015). Convergence, Divergence, and Reconvergence in a Feedforward Network Improves Neural Speed and Accuracy. Neuron 88, 1014–1026. doi:10.1016/j.neuron.2015.10.018.

Jefferis, G. S. X. E., Potter, C. J., Chan, A. M., Marin, E. C., Rohlfing, T., Maurer, C. R., et al. (2007). Comprehensive Maps of Drosophila Higher Olfactory Centers: Spatially Segregated Fruit and Pheromone Representation. Cell 128, 1187–1203. doi:10.1016/j.cell.2007.01.040.

Jinks, A., and Laing, D. G. (1999). A Limit in the Processing of Components in Odour Mixtures. Perception 28, 395–404. doi:10.1068/p2898.

Kadohisa, M., and Wilson, D. A. (2006). Olfactory cortical adaptation facilitates detection of odors against background. J. Neurophysiol. 95, 1888–96. doi:10.1152/jn.00812.2005.

Korner-Nievergelt, F., Roth, T., von Felten, S., Guelat, J., Almasi, B., Korner-Nievergelt, P., et al. (2015). Bayesian data analysis in ecology using linear models with R, BUGS, and Stan. Academic Press.

Krofczik, S., Menzel, R., and Nawrot, M. P. (2009). Rapid odor processing in the honeybee antennal lobe network. Front. Comput. Neurosci. 2, 9. doi:10.3389/neuro.10.009.2008.

Laloi, D., Roger, B., Blight, M. M., Wadhams, L. J., and Pham-Delegue, M. H. (1999). Individual learning ability and complex odor recognition in the honey bee, Apis mellifera L. J. Insect Behav. 12, 585–597. doi:10.1023/A:1020919501871.

Laska, M., and Hudson, R. (1993). Discriminating parts from the whole: determinants of odor mixture perception in squirrel monkeys, Saimiri sciureus. J. Comp. Physiol. A 173, 249–256. doi:10.1007/BF00192984.

Linster, C., Henry, L., Kadohisa, M., and Wilson, D. A. (2007). Synaptic adaptation and odor-background segmentation. Neurobiol. Learn. Mem. 87, 352–360. doi:10.1016/j.nlm.2006.09.011.

Lynn, W. H., Meyer, E. A., Peppiatt, C. E., and Derby, C. D. (1994). Perception of odor mixtures by the spiny lobster Panulirus argus. Chem. Senses 19, 331–47. Available at: http://www.ncbi.nlm.nih.gov/pubmed/7812726 [Accessed: March 19, 2019].

Martelli, C., Carlson, J., and Emonet, T. (2013). Intensity Invariant Dynamics and Odor-Specific Latencies in Olfactory Receptor Neuron Response. J. Neurosci. 33, 6285–6297.

Müller, D., Gerber, B., Hellstern, F., Hammer, M., and Menzel, R. (2000). Sensory preconditioning in honeybees. J. Exp. Biol. 203, 1351–64. Available at: http://www.ncbi.nlm.nih.gov/pubmed/10729283 [Accessed: March 28, 2019].

Murlis, J., Elkinton, J., and Carde, R. (1992). Odor Plumes And How Insects Use Them. Annu. Rev. Entomol. 37, 505–532. doi:10.1146/annurev.ento.37.1.505.

Murlis, J., Willis, M. A., and Carde, R. T. (2000). Spatial and temporal structures of pheromone plumes in fields and forests. Physiol. Entomol. 25, 211–222. doi:10.1046/j.1365-3032.2000.00176.x.

Nagel, K. I., and Wilson, R. I. (2011). Biophysical mechanisms underlying olfactory receptor neuron dynamics. Nat. Neurosci. 14, 208–216. doi:10.1038/nn.2725.

Nikonov, A. A., and Leal, W. S. (2002). Peripheral coding of sex pheromone and a behavioral antagonist in the Japanese beetle, Popillia japonica. J. Chem. Ecol. 28, 1075–89. doi:10.1023/A:1015274104626.

Pavlov, I., Pavlov, P. I., and Anrep, G. V. (1927). Conditioned Reflexes: An Investigation of the Physiological Activity of the Cerebral Cortex., ed. G. Valsievich Anrep London, Oxford University Press.

Pelz, C., Gerber, B., and Menzel, R. (1997). Odorant intensity as a determinant for olfactory conditioning in honeybees: roles in discrimination, overshadowing and memory consolidation. J. Exp. Biol. 200, 837–47. doi:citeulike-article-id:9533426.

Raiser, G., Galizia, C. G. G., and Szyszka, P. (2016). A High-Bandwidth Dual-Channel Olfactory Stimulator for Studying Temporal Sensitivity of Olfactory Processing. Chem. Senses 42, bjw114. doi:10.1093/chemse/bjw114.

Reinhard, J., Sinclair, M., Srinivasan, M. V., and Claudianos, C. (2010). Honeybees Learn Odour Mixtures via a Selection of Key Odorants. PLoS One 5, e9110. doi:10.1371/journal.pone.0009110.

Riffell, J. A., Shlizerman, E., Sanders, E., Abrell, L., Medina, B., Hinterwirth, A. J., et al. (2014). Sensory biology. Flower discrimination by pollinators in a dynamic chemical environment. Science 344, 1515–8. doi:10.1126/science.1251041.

Rokni, D., Hemmelder, V., Kapoor, V., and Murthy, V. N. (2014). An olfactory cocktail party: figure-ground segregation of odorants in rodents. Nat. Neurosci. 17, 1225–1232. doi:10.1038/nn.3775.

Roussel, E., Carcaud, J., Combe, M., Giurfa, M., and Sandoz, J.-C. (2014). Olfactory Coding in the Honeybee Lateral Horn. Curr. Biol. 24, 561–567. doi:10.1016/j.cub.2014.01.063.

Sachse, S., and Galizia, C. G. (2002). Role of inhibition for temporal and spatial odor representation in olfactory output neurons: a calcium imaging study. J. Neurophysiol. 87, 1106–1117. doi:11826074.

Saha, D., Leong, K., Li, C., Peterson, S., Siegel, G., and Raman, B. (2013). A spatiotemporal coding mechanism for background-invariant odor recognition. Nat. Neurosci. 16, 1830–9. doi:10.1038/nn.3570.

Sehdev, A., Mohammed, Y. G., Triphan, T., and Szyszka, P. (2019). Olfactory Object Recognition Based on Fine-Scale Stimulus Timing in Drosophila. iScience 13, 113–124. doi:10.1016/j.isci.2019.02.014.

Smith, B. (1998). Analysis of interaction in binary odorant mixtures. Physiol. Behav. 65, 397–407.

Smith, B. H., and Cobey, S. (1994). The olfactory memory of the honeybee Apis mellifera. II. Blocking between odorants in binary mixtures. J. Exp. Biol. 195, 91–108. Available at: http://www.ncbi.nlm.nih.gov/pubmed/7964421 [Accessed: May 14, 2017].

Staubli, U., Fraser, D., Faraday, R., and Lynch, G. (1987). Olfaction and the “data” memory system in rats. Behav. Neurosci. 101, 757–65. Available at: http://www.ncbi.nlm.nih.gov/pubmed/3426792 [Accessed: March 19, 2019].

Stevenson, R. J., and Wilson, D. A. (2007). Odour perception: An object-recognition approach. Perception 36, 1821–1833. doi:10.1068/p5563.

Stopfer, M., and Laurent, G. (1999). Short-term memory in olfactory network dynamics. Nature 402, 664–668. doi:10.1038/45244.

Strube-Bloss, M. F., Nawrot, M. P., and Menzel, R. (2011). Mushroom Body Output Neurons Encode Odor-Reward Associations. J. Neurosci. 31, 3129–3140. doi:10.1523/JNEUROSCI.2583-10.2011.

Strutz, A., Soelter, J., Baschwitz, A., Farhan, A., Grabe, V., Rybak, J., et al. (2014). Decoding odor quality and intensity in the Drosophila brain. Elife 3, e04147. doi:10.7554/eLife.04147.

Szyszka, P., Ditzen, M., Galkin, A., Galizia, C. G. G., and Menzel, R. (2005). Sparsening and Temporal Sharpening of Olfactory Representations in the Honeybee Mushroom Bodies. J. Neurophysiol. 94, 3303–3313. doi:10.1152/jn.00397.2005.

Szyszka, P., Stierle, J. S., Biergans, S., and Galizia, C. G. (2012). The Speed of Smell: Odor-Object Segregation within Milliseconds. PLoS One 7, e36096. doi:10.1371/journal.pone.0036096.

Weissburg, M., Atkins, L., Berkenkamp, K., and Mankin, D. (2012). Dine or dash? Turbulence inhibits blue crab navigation in attractive-aversive odor plumes by altering signal structure encoded by the olfactory pathway. J. Exp. Biol. 215, 4175–82. doi:10.1242/jeb.077255.

Wilson, D. A. (1998). Synaptic Correlates of Odor Habituation in the Rat Anterior Piriform Cortex. J. Neurophysiol. 83, 139–145. doi:10.1152/jn.2000.83.1.139.

Wright, G. A., and Smith, B. H. (2004). Variation in complex olfactory stimuli and its influence on odour recognition. Proc. R. Soc. B Biol. Sci. 271, 147–152. doi:10.1098/rspb.2003.2590.

